# Prediction of bacterial protein-compound interactions with only positive samples

**DOI:** 10.1101/2025.07.18.665643

**Authors:** Ki-Hwa Kim, Avinash Yaganapu, Sai Kosaraju, Aashish Bhatt, Yun Lyna Luo, Sai Phani Parsa, Juyeon Park, Hyun Lee, Jun Hyuck Lee, Tae-Jin Oh, Mingon Kang

**Affiliations:** Genome-based BioIT Convergence Institute, Asan, Republic of Korea; Department of Computer Science, University of Nevada, Las Vegas, Las Vegas, NV, USA; Department of Computer Science, California State Polytechnic University, Pomona, CA, USA; Department of Biotechnology and Pharmaceutical Sciences, Western University of Health Sciences, Pomona, CA, USA; Department of Computer Science, Sun Moon University, Asan, Republic of Korea; Division of Life Sciences, Korea Polar Research Institute, Incheon, Republic of Korea; Department of AI Biomedical Engineering, Graduate School, Sun Moon University, Asan, Republic of Korea; Department of Pharmaceutical Engineering and Biotechnology, Sun Moon University, Asan, Republic of Korea

**Keywords:** Compound-protein interaction, bacterial CYP, biocatalyst, deep learning, positive-unlabeled learning

## Abstract

Prediction of Compound-Protein Interactions (CPI) in bacteria is crucial to advance various pharmaceutical and chemical engineering fields, including bio-catalysis, drug discovery, and industrial processing. However, current CPI models cannot be applied for bacterial CPI prediction due to the lack of curated negative interaction samples. This paper introduces a novel Positive-Unlabeled (PU) learning framework, named BIN-PU, to address this limitation. BIN-PU generates pseudo positive and negative labels from known positive interaction data, enabling effective training of deep learning models for CPI prediction. We also propose a weighted positive loss function that weights to truly positive samples. We have validated BIN-PU with multiple CPI backbone models, comparing the performance with the existing PU model using bacterial cytochrome P450 (CYP) data. Extensive experiments demonstrate the superiority of BIN-PU over the benchmark model in predicting CPIs with only truly positive samples. Furthermore, we have validated BIN-PU on additional bacterial proteins obtained from literature review, human CYP datasets, and uncurated data for its reproducibility. We have also validated the CPI prediction for the uncurated CYP data with biological and biophysical experiments. BIN-PU represents a significant advancement in CPI prediction for bacterial proteins, opening new possibilities for improving predictive models in related biological interaction tasks.

## 1 Introduction

Accurate prediction of Compound-Protein Interactions (CPI) in bacteria is essential in discovering diverse chemical properties and understanding intermediate mechanisms of biosynthesis and biodegradation [1, 2]. Bacterial enzymes act as bio-catalysts that react with compounds and are widely utilized in sectors including agriculture, chemical industry, food, textiles, pharmaceuticals, cosmetics, and bioremediation [1, 3]. Bacterial proteins metabolize a wide spectrum of endogenous and exogenous compounds, enhancing the efficiency of biocatalyst processes that selectively interact with bioactive compounds [4, 5]. However, characterizing biological interactions between proteins and compounds remains extremely challenging due to the complexity of protein networks, heterogeneity in protein behavior across environments, and high substrate selectivity [6–8].

Deep learning-based CPI models have been explored using rich reference databases, particularly focusing on human proteins and compounds. These models learn relevant features directly from protein and compound data, enhancing their capability to discover complex interactions—unlike traditional machine learning approaches that rely on predefined features, such as kernel-based SVMs [9–12], graph data mining [13], and tree-based classifiers [14]. Moreover, non-linear relationships and hierarchical representations are effectively captured by deep learning models, which are crucial for understanding CPIs. Current state-of-the-art deep learning CPI models utilize Graph Neural Networks (GNNs), Convolutional Neural Networks (CNNs), and Transformers. For instance, CPIprediction encodes protein sequences using CNN and compounds using GNN, followed by feature integration to predict their interactions [15]. TransformerCPI employs a transformer-based framework with an encoder-decoder structure to extract low-level representations of proteins and compounds [16]. SSNet incorporates protein secondary structure databases along with CNN and GNN-based representations for both proteins and compounds [17]. However, all these CPI models require both positive (i.e., known interactions) and negative (i.e., known non-interactions) samples for training, which limits their applicability to bacterial CPI prediction.

Deep learning models for bacterial CPI prediction have seldom been developed, primarily due to the lack of databases containing known non-interactions. In microorganisms, the extremely limited availability of negative samples presents a significant barrier to utilizing current state-of-the-art models that depend on both positive and negative interactions. Several factors contribute to this challenge. First, the complexity and diversity of bacterial species hinder the identification of true negative samples, as interaction likelihood varies across species and environmental contexts. Moreover, experimental designs involving heterologous expression of bacterial proteins may not accurately represent their native environments [18]. Consequently, the absence of inactive samples creates a substantial gap, limiting the development of effective CPI prediction models for bacterial systems.

Positive-Unlabeled (PU) learning presents a promising solution by enabling model training using only positive samples. PU learning typically extends the training set by inferring pseudo labels for likely positive and negative examples derived solely from truly positive data [19]. Generative Adversarial Networks (GANs) have been leveraged in PU settings, where a generator produces synthetic negatives and a discriminator distinguishes between generated and real positives [20, 21]. Alternatively, sample-wise weighting schemes, in which positive samples are assigned greater weights than unlabeled data, have been proposed to improve classification performance by penalizing misclassification and ensuring consistency through resampling [22, 23].

In this study, we propose an effective PU learning framework, BIN-PU (BINning strategy for PU), to predict interactions between bacterial proteins and compounds. BIN-PU generates reliable pseudo positive and negative labels from truly positive data and builds a classifier with a weighted positive loss function to predict CPIs (Fig. 1). We evaluated BIN-PU’s robust predictive performance in multiple settings. First, we assessed BIN-PU coupled with CPI backbone models on the cross-validation dataset, compared to the existing PU strategy. Second, we validated BIN-PU using independent bacterial CYP from literature and curated human enzymes. Finally, we applied BIN-PU to uncurated bacterial CYPs to explore novel enzyme–substrate interactions. The predicted interactions were experimentally verified through biological assays (HPLC) and biophysical analysis. The details of the BIN-PU framework and its implementation are elucidated in Methods. In summary, our main contributions are: (1) BIN-PU predicts potential interactions between compounds and bacterial proteins using only truly positive samples; (2) Extensive experiments coupled with CPI backbone models were conducted to validate the effectiveness and advantages of the proposed PU strategy, compared to the current state-of-the-art method PUCPI; (3) We further validated the model using bacterial CYP and substrate data through laboratory assays, integrating both biological validation and molecular docking studies. To the best of our knowledge, this is the first method to predict CPIs for bacterial enzymes using only truly positive dataset.

**Fig. 1.**
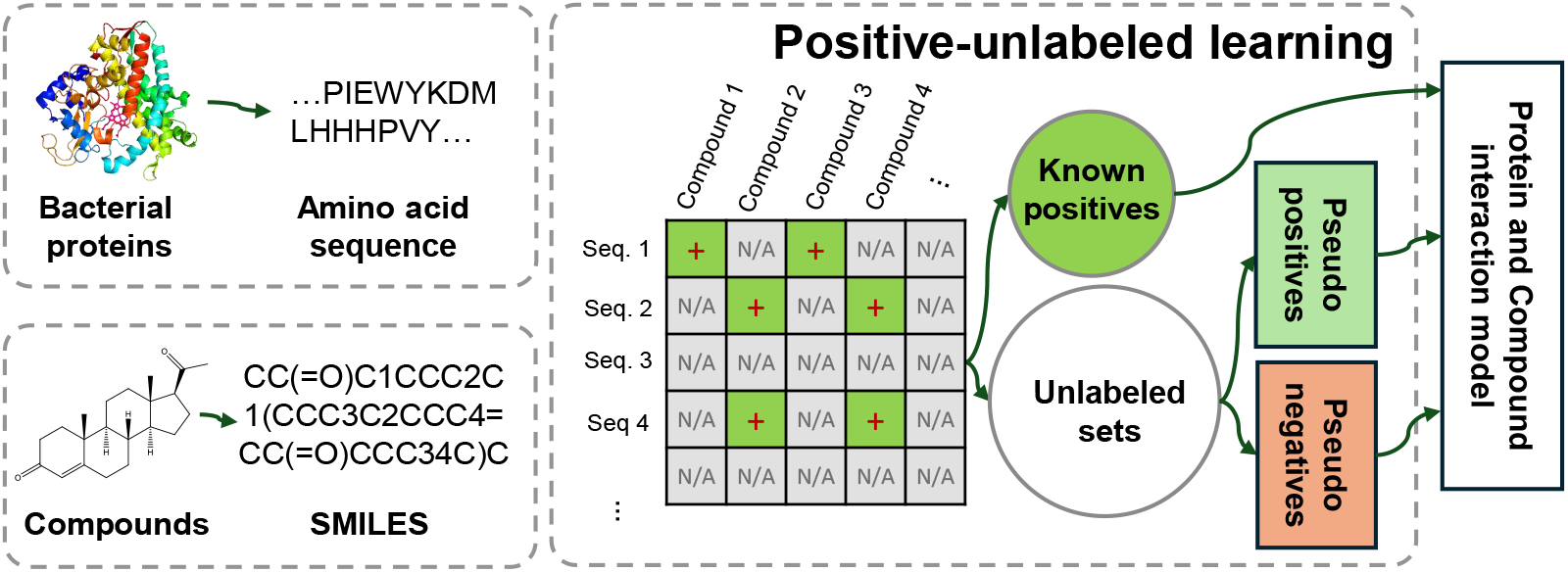
Overview of the study. The proposed PU learning strategy trains deep learning models to predict interactions between bacterial proteins and compounds using only known interaction databases by generating pseudo positives and negatives from unlabeled datasets.

## 2 Results

### 2.1 Dataset for model development

In this study, we focus on bacterial CYP, which is a well-known monooxygenase that catalyzes secondary metabolism, such as biosynthesis of steroid hormones. CYP consists of a unique protein structure that contains heme molecules. Heme molecules induce modification of the backbone or side chain of the compound through redox reactions. CYP is a candidate that may interact with diverse compounds for unknown metabolism, due to its characteristics, such as regioselectivity, stereoselectivity, and a broad substrate spectrum. Furthermore, CYP exists in all living things, as it has been isolated from mammals, bacteria, and even viruses.

We collected labeled bacteria data from the two databases of Protein Data Bank (PDB) and UniProtKB/Swiss-Prot (Swiss-Prot). The two databases include proteins, compounds, and their interactions, where proteins and compounds are provided as sequences of amino acids and SMILE strings converted from InChIKey, respectively. We searched the PDB and Swiss-Prot databases using the keywords ‘Cytochrome P450’ and ‘bacteria’. To enhance the quality of the datasets, we manually examined the dataset. We removed non-CYP proteins and irrelevant proteins and ligands (Supplementary Note S1, Supplementary Table S1 and S2). In PDB and Swiss-Prot, we excluded non-binding ligands (e.g., ionic compounds, small molecules, and linkers other than the substrate) and HEM. To label the interactions, we referred to the column of ‘unique ligands’ for each enzyme entry in PDB and ‘catalytic activity’ in Swiss-Prot as truly positive samples. We finally obtained 500 truly positive entries of unique compounds (*N* = 291) and proteins (*N* = 241). Then, we generated unlabeled data by considering the pairwise combinations. The unlabeled data excluded the known interactions (i.e., positive samples), resulting in 69,631 entries that include both likely-positive and negative samples.

### 2.2 Assessment of the predictive performance with cross validation

We evaluated the performance of our proposed BIN-PU compared with PUCPI. We performed two assessments in this cross-validation experiment: (1) assessment of pseudo labels and (2) evaluation of the performance in a fully supervised learning manner on the test data. First, we generated the pseudo labels from truly positive and unlabeled data and assessed the quality of the pseudo labels on validation data. Then, we trained BIN-PU using the truly positive and reliable pseudo labeled samples coupled with CPI backbone models in a fully supervised manner. We compared the performance of BIN-PU with PUCPI using test data. We considered SSNet [17], CPIprediction [15], and TransformerCPI [16] as backbone CPI models for both BIN-PU and PUCPI. Detailed implementation and tuning of these models are provided in the supplementary materials (Supplementary Note S2–S3).

For the entire experiment, we randomly split the truly positive samples into 60% for training (*N* = 300), 20% for validation (*N* = 100), and 20% for testing (*N* = 100). Then, BIN-PU generated pseudo labels from the unlabeled data of the training. The generated pseudo labeled samples were again split into 60%, 20%, and 20% to aggregate with the truly positive samples in the training, validation, and test group, respectively. The final sample sizes were (*N* = 20,178) for training, (*N* = 6,726) for validation, and (*N* = 6,726) for test on average. We repeated all of the experiments ten times to ensure reproducibility.

For the assessment of the quality of the pseudo labels, we computed Spy Capture Rates (SCR). We calculated SCR by counting the number of positive predictions among the spy samples in the multiple bins. The SCR scores for BIN-PU coupled with SSNet, CPIprediction, and TransformerCPI were 0.82 ±0.04, 0.91 ±0.03, and 0.93 ±0.02, respectively (Supplementary Table S3). BIN-PU with TransformerCPI achieved the highest SCR, which implies that 93% of the spy samples are correctly identified by BIN-PU as positive in the unlabeled data. Note that PUCPI does not generate pseudo labels, and SCR was not compared.

For the assessment of the predictive performance, we computed F1-scores to compare the performance between BIN-PU and PUCPI. For BIN-PU, we empirically optimized hyper-parameters using the validation data. Learning rate (i.e., 1e − 4), weight decay (i.e., 1e − 5), and bin size (20) were optimized to maximize SCR, whereas the optimal weight parameter (*λ*) in (4) and the threshold for the final discriminant function were obtained by maximizing F1-scores in the validation data (Supplementary Fig. S1 and Supplementary Table S4). For PUCPI, we considered the same CPI backbone models for fair comparison with BIN-PU along with biased SVM that the original PUCPI study used. For PUCPI, we empirically determined the optimal weight for positive samples across all three CPI backbone models, as well as in the biased-SVM model. In PUCPI coupled with a biased-SVM, the threshold for the final discriminant function was optimized to maximize F1-scores with the validation data, using the pseudo labels generated by BIN-PU coupled with TransformerCPI. The training procedures and hyperparameter tuning of all three CPI backbone models and the biased-SVM model were provided in the supplementary (Supplementary Note S2–S3).

The F1-scores of both BIN-PU and PUCPI coupled with the CPI backbone models are illustrated in Fig. 2A. BIN-PU consistently outperformed PUCPI across all of the three backbone models. Specifically, BIN-PU with TransformerCPI achieved the highest F1-score of 0.8216±0.0012, compared to PUCPI of 0.7527±0.0016, reflecting a significant performance improvement of approximately 7%. The improvement was statistically validated by the Wilcoxon rank-sum test (*p* < 0.05). BIN-PU also showed notable improvements with the other benchmark models of CPIprediction and SSNet, approximately 7% and 9%, with averaged F1-scores of 0.7372±0.0030 and 0.7046±0.0086, whereas PUCPI’s were 0.6686±0.0034 and 0.6137±0.0093 (Wilcoxon rank-sum test *p*-value < 0.05).

**Fig. 2.**
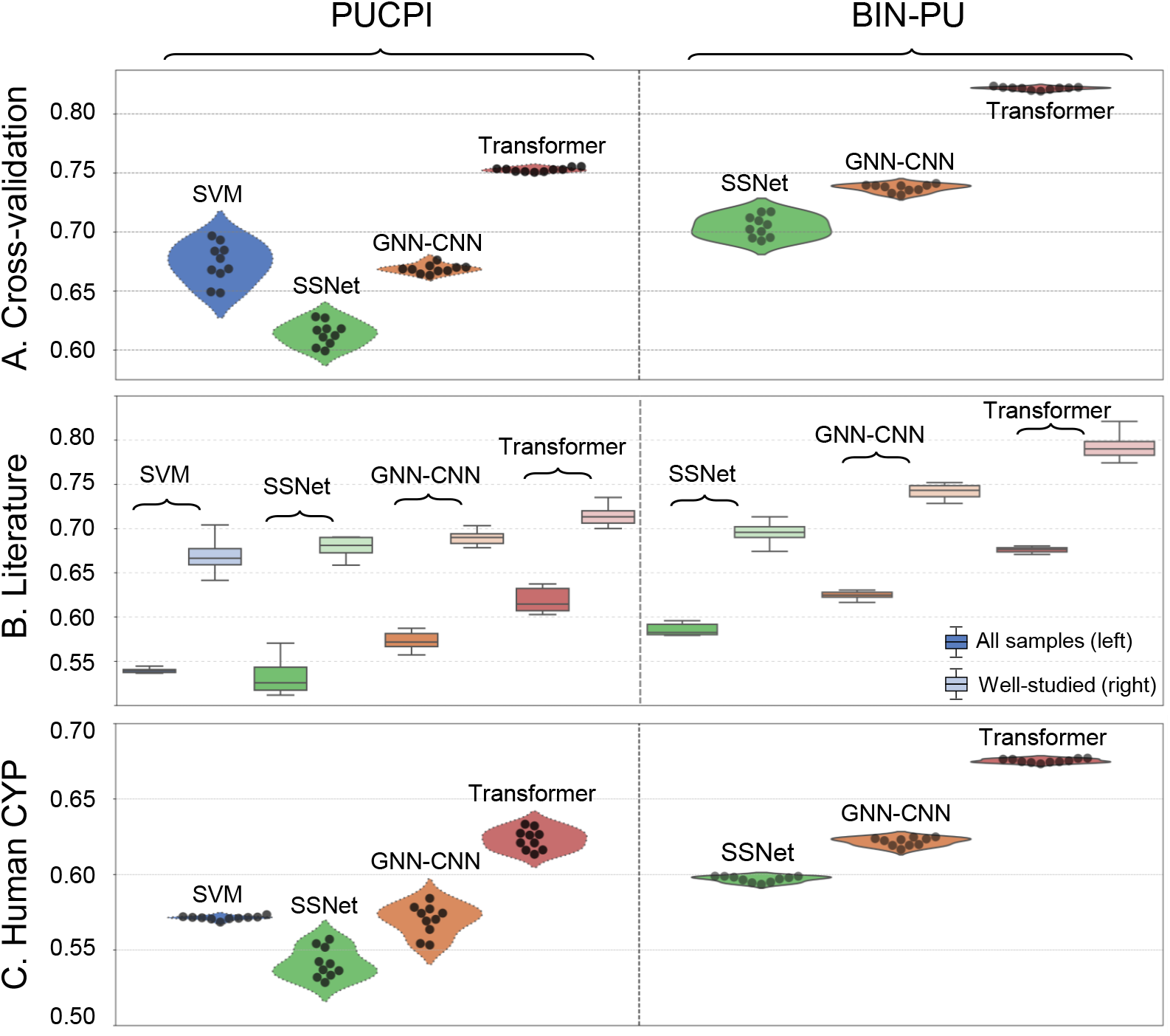
The performance comparison between BIN-PU and PUCPI coupled with CPI prediction models on F1 scores. F1 scores (A) in the cross-validation experiments, (B) with additional entire bacterial samples (*N* = 129) from the literature (on the left side) and only 41 samples with 30 well-studied substrates among them in the right side, and (C) with human CYP of both positive and negative samples. The BIN-PU was integrated with SSNet-, GNN-CNN-, and Transformer-based CPI models, while the existing PU strategy (PUCPI) is applied to these models as well as biased-SVM.

### 2.3 External validation with literature data in bacterial CYP

We evaluated the performance of BIN-PU using additional bacterial samples manually collected from the literature. We have identified bacterial CYP samples (Supplementary Table S5). We searched the literature published from 2000 to 2022 using the keyword, “microbial CYP family” in PubMed, limiting it to the pairs of five CYPs and 98 substrates. Finally, we collected 80 positive and 49 negative CPI samples. We confirmed that the external literature validation data were not included in the crossvalidation data. For the performance assessment, we computed F1-scores using the optimal BIN-PU and PUCPI models, which were trained with the cross-validation data (Fig. 2B). BIN-PU showed significantly improved performance with all the CPI backbone models, compared to PUCPI. BIN-PU with TransformerCPI achieved the highest F1-score of 0.6788±0.0033 (Wilcoxon rank-sum test *p*-value < 0.05). PUCPI with TransformerCPI was 0.6374±0.0127. Similarly, BIN-PU with CPIprediction and SSNet obtained the higher F1-scores of 0.6303±0.0039 and 0.5957±0.0109, compared to PUCPI’s 0.5848±0.0097 and 0.5528±0.0181 (Wilcoxon rank-sum test *p*-value < 0.05). The overall low accuracy was mainly due to limited interaction information in the diverse substrates in the training dataset. Thus, we narrowed down the examination, focusing on 30 well-studied substrates with simple structures. When we considered the well-studied samples (*N* = 41), the accuracy significantly improved to 79.3% by BIN-PU with TransformerCPI. This may imply the importance of comprehensive and high-quality training data for enhancing predictive performance and reliability.

### 2.4 External validation with curated human CYP data

We further evaluated the performance using human CYP data, in which negative ground truths are available, for the indirect assessment. The study was with the assumption that fundamental patterns of CYP proteins are shared across the species. We obtained the data from Integrated Protein-Ligand Interaction Database (IPLID), which comprises 17,273 unique substrates and 47 unique CYP proteins [24]. The human CYP dataset contained 25,045 positive samples and 37,277 negative samples. We applied the final models, which were trained with the cross-validation of bacteria CYP data, to the human data. Fig. 3C depicts the F1-scores on the human CYP dataset. BIN-PU also outperformed PUCPI. BIN-PU with TransformerCPI achieved the highest F1-score of 0.6762±0.0012, compared to PUCPI of 0.6332±0.0065 (Wilcoxon rank-sum test *p*-value < 0.05). Similarly, BIN-PU with CPIprediction and SSNet showed higher F1-scores than PUCPI, with F1-scores of 0.6229±0.0026 and 0.5978±0.0018 compared to PUCPI’s 0.5742±0.0096 and 0.5570±0.0094, respectively (Wilcoxon rank-sum test *p*-value < 0.05).

**Fig. 3.**
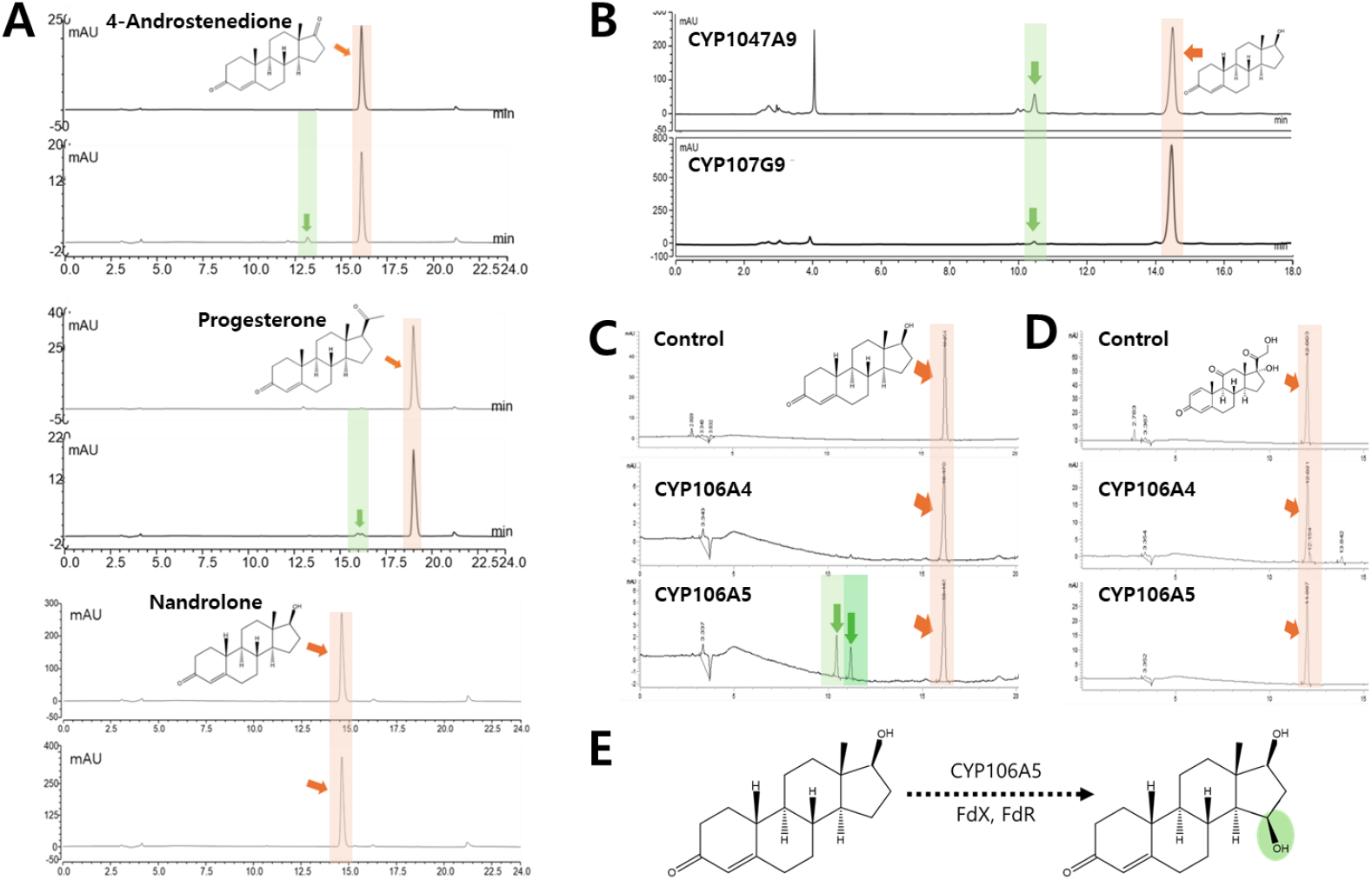
HPLC chromatogram of the *in vitro* conversion of steroid by CYPs. (A) HPLC analysis of CYP154C9 and three steroids (4-Androstenedione, progesterone, and nandrolone) (Exp.1–3); (B) Activity analysis of CYP1047A9 and CYP107G9 with nandrolone (Exp.4 and 5); (C) Comparison of nandrolone activities between CYP106A4 and CYP106A5 (Exp.6 and 7); (D) Activity analysis of CYP106A4 and CYP106A5 with prednisone (Exp.8 and 9). Orange arrows and boxes indicate sub-strate peaks, and green arrows and boxes indicate products; (E) Structure of the predicted product in CYP106A5.

### 2.5 Validation with uncurated bacterial CYP data

We applied the proposed models to uncurated bacterial CYP proteins with various substrates to identify their potential interactions and validated them with biological and biophysical experiments, as it is a typical practice in genome studies. Note that the previous experiments validated the models using well-established curated data only. In this experiment, we explored five CYP proteins of CYP154C9, CYP1047A9, CYP107G9, CYP106A4, and CYP106A5, which are potential candidates of novel steroid hydroxylase. For substrates, we considered four steroids of androstenedione, progesterone, nandrolone, and prednisone, known as representative substrates for CYP due to their well-established interactions with CYP. Along with the five CYP proteins and four steroids, we mainly examined their interactions focusing on: (1) the interactions of a CYP protein (CYP154C9) with the three steroids (Exp. 1–3 in Table 1), (2) the interactions of a steroid (nandrolone) with the five CYP proteins (Exp. 3–7 in Table 1), and (3) the interactions of a steroid (prednisone) with the two CYP proteins (Exp. 8 and 9 in Table 1).

**Table 1.**
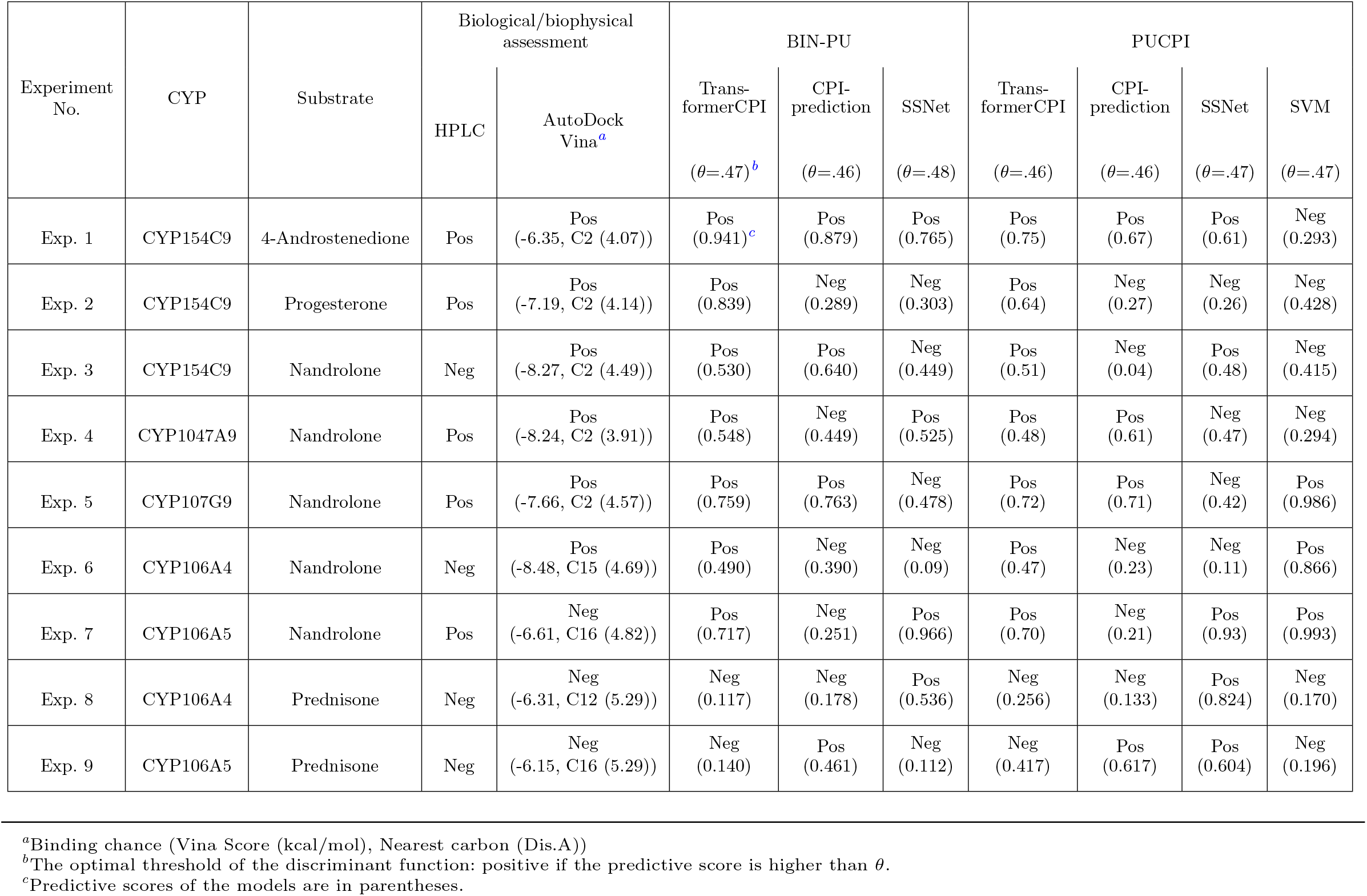
CPI scores of the five uncurated CYP and the three steroids. The reactions were validated through biological (HPLC) and biophysical (AutoDock Vina) experiments.

We presumed their potential reactions by conducting both biological and biophysical experiments. To biologically assess the potential reactions, we cultured, purified, and conducted in vitro activity assays. We obtained bacterial strains of Streptomyces alboniger, Streptomyces sp., and Actinomycete sp. from the Korean Culture Center of Microorganisms (KCCM) and the American Type Culture Collection (ATCC) (Supplementary Table S6). Additionally, we obtained a Paenibacillus sp. strain isolated from Antarctica to expand the diversity of microbial sources for enzyme characterization. We identified the five CYP genes in the microorganisms based on gene sequence identity, including the conserved signature heme-binding domain (FXXGX(H/R)XCXG), and the CYP enzyme names were assigned by Dr. David Nelson [25]. We amplified the target genes using Polymerase Chain Reaction (PCR), which were subsequently cloned into pET-28a(+) or pET-32a(+) expression vectors and expressed in recombinant Escherichia coli C41 (DE3) strains. The target genes were transcribed and translated into proteins, followed by purification using a Lac operon-based overexpression system and a His-tag method to obtain high-purity protein. We performed in vitro enzymatic activity assays using the purified proteins, and validated the reactions through High-Performance Liquid Chromatography (HPLC) (Supplementary Note S4, Fig. 3). In the HPLC analysis, inactivated CYP served as a control, allowing the detection of novel peaks corresponding to the products formed from the activity reaction.

The HPLC analyses estimated five reactions (Exp. 1–2, 4–5, and 7) and four non-reactions (Exp. 3, 6, 8, and 9). New products in the HPLC analysis (peaks in green, Fig. 3A) indicate reactions of CYP154C9 with androstenedione and progesterone (Exp. 1–2), while no evidence was shown for the interaction with nandrolone (Exp. 3). Similarly, new products, generated by reaction of nandrolone on CYP1047A9 and CYP107G9, were observed in the HPLC analysis (Fig. 3B, Exp. 4–5), indicating their reactions. CYP106A4 showed no interaction with nandrolone (Exp. 6), whereas CYP106A5 showed reactions with nandrolone (Exp. 7), although they have approximately 55% sequence similarity (Fig. 3C). CYP106A4 and CYP106A5 did not react with prednisone (Fig. 3D, Exp. 8–9). Notably, HPLC of CYP106A5 revealed two distinct product peaks, with one matching the retention time of a 15-beta-hydroxylated nandrolone reported for BaCYP106A2 (Fig. 3E, Supplementary Fig. S2) [26].

We also conducted biophysical experiments using homology modeling and molecular docking to further assess potential protein–ligand interactions. As the three-dimensional structures of the five CYP strains were not available in the Protein Data Bank, homology models of the CYP proteins were generated using the Chai-1 server [27]. The substrate compounds were retrieved from the PubChem database. Binding affinity scores and distances between the heme group and the nearest carbon atom by AutoDock Vina (version 1.2.3) and chimeraX, and the protein–compound interactions were measured by using the MOE software (the details are in Supplementary Note S5 and Supplementary Table S7) [28]. The results of this analysis are presented in Table 1.

The biophysical assessments revealed five strong binding affinities in Exp. 2–6 and comparatively weaker interactions in Exp. 1 and 7–9 (Table 1). Binding energies in molecular docking are typically expressed as negative values, where lower (more negative) scores reflect stronger binding affinities. In Exp. 1–3, CYP154C9 was docked with three substrates, and the docked poses indicated potential reactions at carbon 2. It exhibited strong binding scores of − .19 kcal/mol with progesterone in Exp. 2 and − 8.27 kcal/mol with nandrolone in Exp. 3, whereas a weaker affinity of − 6.35 kcal/-mol was observed with 4-androstenedione in Exp. 1. This observation aligns with the known catalytic function of CYP154C2, another member of the CYP154C family, which has been reported to catalyze hydroxylation at the same carbon position 2*α* [29]. In Exp. 4–5, CYP1047A9 and CYP107G9 showed strong binding affinities of − 8.24 and − 7.66 kcal/mol, respectively, implying possible reactivity at carbon 2. However, these novel CYPs remain uncharacterized due to limited research on similar family members. In Exp. 6–9, docking analysis of CYP106A4 and CYP106A5 revealed two substrate binding poses. The interaction between CYP106A4 and nandrolone in Exp. 6 exhibited the strongest affinity (− 8.48 kcal/mol), suggesting potential hydroxylation at carbon 15. This is supported by studies on CYP106A2, CYP106A6, and other CYP106A family members known to catalyze 15*β*-hydroxylation of steroid substrates [30, 31]. Comparatively weaker binding energies were observed in Exp. 7–9 (− 6.61, − 6.31, and − 6.15 kcal/mol, respectively). The low affinity observed in prednisone interactions (Exp. 8–9) may be due to steric hindrance from the complex branching at position C17, impeding effective binding to the active site.

Although molecular modeling analyses suggested the possibility of binding with substrate in Exp. 3 and 6, no product formation was observed in the biological assays. Biological enzymatic experiments are highly sensitive to biochemical factors (e.g., substrate hydrophobicity, electron donor systems) and environmental conditions (e.g., temperature, pH, and buffer composition). We conducted experiments following the optimal conditions for each CYP. CYP154C9 was tested using iodobenzene, which is a known effective electron transfer system for the CYP154C family (Exp. 3) [32]. Similarly, CYP106A4 was examined under the Fdx–FdR electron transfer system, which has demonstrated catalytic activity in CYP106A1, BaCYP106A2, and BaCYP106A6 (Exp. 6) [ref]. Nevertheless, the novel CYPs used in this study exhibited low expression levels and weak catalytic reactivity, which could potentially be improved through further optimization of the experimental conditions.

Finally, based on the biological and biophysical assessment, we presumably concluded four positive interactions in Exp. 1, 2, 4, and 5, three weakly possible interactions in Exp. 3, 6, and 7, and two non-interactions in Exp. 8 and 9. For the weakly possible interactions, Exp. 3 and 6 were shown as non-reaction in biological assays, whereas Exp. 7 was positive. Therefore, the product in Exp. 7, which was confirmed by the biological experiment, suggests the possibility of a novel product hydroxylated at carbon 16, as supported by the biophysical analysis. In contrast, Exp. 3 and 6, which yielded no product in biological assays, were further analyzed using MOE to investigate interactions between amino acids for more detailed insights. Further analysis of the docking interactions revealed potential binding patterns for each steroid compound (Fig. 4 and Supplementary Fig. S3). Exp. 6 exhibited the strongest binding affinity by AutoDock among the nine experiments, where the ligand formed a backbone acceptor hydrogen bond with amino acid residues Ala293, Lys294, and Pro397. Exp. 3, showing the second strongest binding affinity by AutoDock, observed a hydrogen bond with the residue Thr282. These docking results offered structural insights of the binding modes and potential reactivity of the steroid compounds with the tested five CYP enzymes (Supplementary Fig. S3). Interestingly, nandrolone exhibited favorable binding energies and interactions with both CYP154C9 and CYP106A4 (Exp. 3 and 6), suggesting its potential as a promising substrate for steroid hydroxylation.

**Fig. 4.**
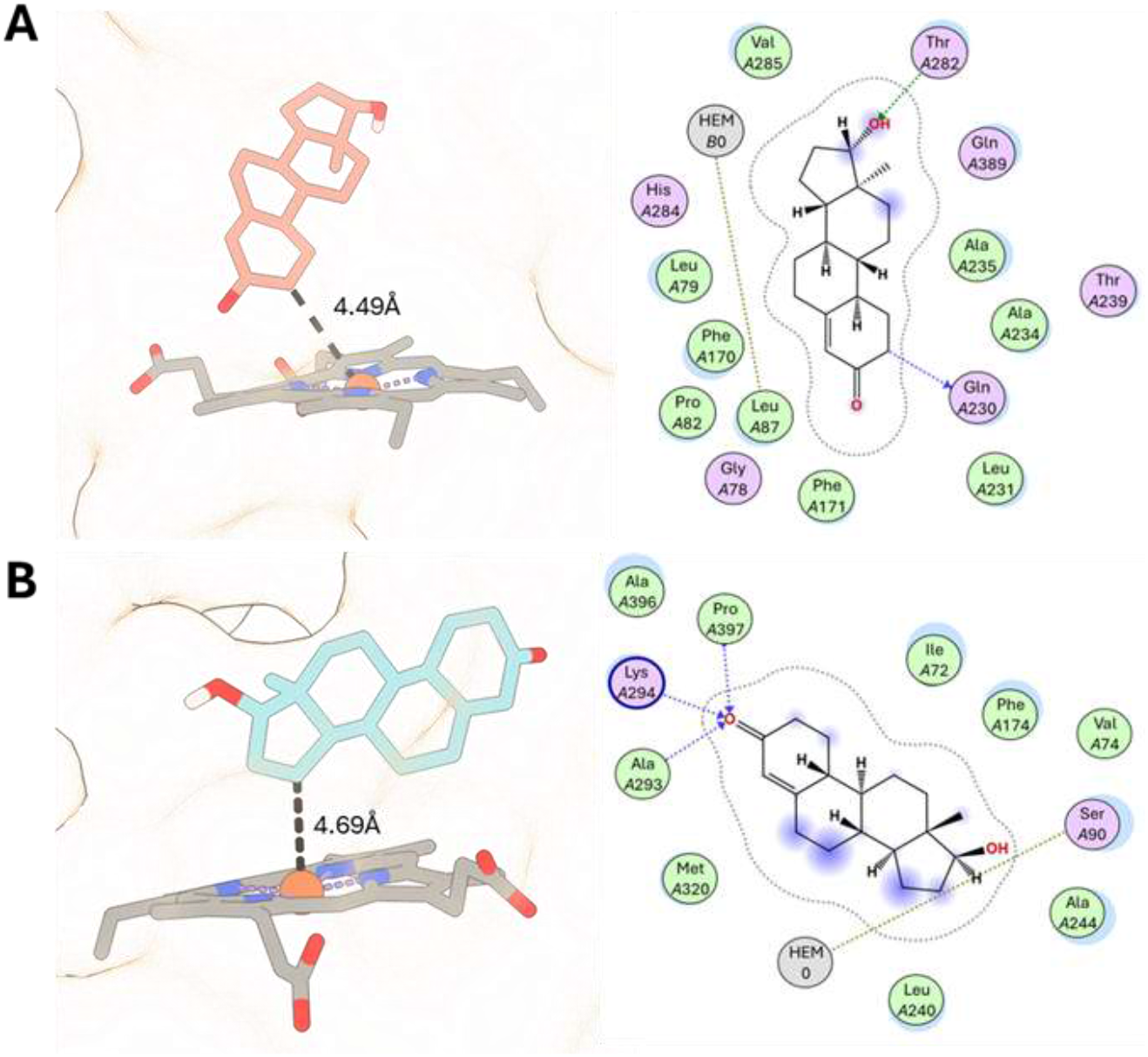
Binding mode in active site and 2D molecular interaction diagrams of the best docked compounds. (A) Active site of CYP154C9 with nandrolone. (B) Binding pose of CYP106A4 protein with nandrolone and their interaction.

We applied BIN-PU and PUCPI to predict the potential interactions between the CYPs and steroids (Table 1). For the evaluation, we mainly considered BIN-PU with TransformerCPI, which produced the best model in the previous experiments, while the predictive results with other models are also shown. BIN-PU with TransformerCPI identified seven potential interactions in Exp. 1–7 and two non-interactions in Exp. 8–9 (Table 1). In the positive predictions in Exp. 1–7, the highest prediction score was observed in Exp. 1 for the CYP154C9-androstenedione reaction, with a predictive score of 0.941, followed by Exp. 2 (0.839), Exp. 5 (0.759), Exp. 7 (0.717), and Exp. 4 (0.548). On the other hand, Exp. 3 and 6 showed weak predictive scores of 0.530 and 0.490, respectively, by BIN-PU with TransformerCPI. In Exp. 8 and 9, BIN-PU with TransformerCPI predicted no interaction with scores of 0.117 and 0.140, respectively.

## 3 Conclusions

We introduced a novel positive-unlabeled learning strategy (BIN-PU) for CPI prediction, especially where curated negative samples are not available. BIN-PU integrates unlabeled samples with truly positive samples, mitigating the challenge of lack of negative samples. We conducted intensive experiments with various experimental settings and bacterial CYP datasets, and the experiments demonstrated the effectiveness of our approach across various CPI backbone architectures with a significantly improved predictive performance than PUCPI. We also applied it to uncurated CYP and substrates, and the potential reaction predictions were validated with biological and biophysical experiments.

In this study, we considered both activators and inhibitors as reactions due to the limited availability of labeled data of inhibitors and activators for the interactions in this study. Nevertheless, CPI prediction using BIN-PU can be an effective strategy in identifying potential substrates in early-stage biocatalytic applications or virtual screening processes. The limitation could be further tackled by integrating additional biochemical data or training with labeled datasets of activators and inhibitors available in the future.

We demonstrates potential deep learning-based applications using only positive interaction labels by mainly assessing it with bacterial CYP proteins and related substrates. Future work for diverse ranges of bacterial proteins and substrates will be required to generally apply it for bacteria protein-substrate interactions prediction. Eventually, prediction of novel bacterial enzyme-substrate interactions may serve as a foundation for pharmaceutical applications and enhance biocatalysis and drug discovery.

## 4 Methods

### 4.1 Current PU learning strategy for CPI

The only PU learning strategy in CPI is PUCPI [23], although PUCPI mainly investigates human data. PUCPI trains a binary classifier, where unlabeled data is considered as negative with the assumption that the unlabeled data primarily contains negative examples. PUCPI generates unlabeled samples derived from the compound-protein pairs of truly positive samples and determines the decision boundary by assigning weights to truly positive samples. The PUCPI study used biased SVM to implement the class weights [23]. However, PUCPI has the following limitations: (1) PUCPI has a strict assumption that the majority of the unlabeled data is negative, which results in the decision boundary biased from the unreliable negative samples; (2) PUCPI assigns more weights to positive instances, but it faces challenges in effectively handling imbalanced data; (3) non-availability of reliable negative samples make the evaluation difficult; and (4) PUCPI was validated by using limited positive samples only with the external data, making it difficult to assess the false negative rate or the reliability of the model’s negative predictions.

### 4.2 The proposed PU learning strategy

We introduce a novel PU learning strategy, BIN-PU, that tackles the limitations of PUCPI. The overall pipeline of BIN-PU consists of the steps: (1) generating unlabeled data, (2) generating pseudo labels of interactions between proteins and compounds only from positive data (the key function of BIN-PU), and (3) training supervised backbone CPI prediction models using the pseudo positive and negative labels (Fig. 5). Unlabeled data is generated from the pairwise combinations between unique proteins and compounds in truly positive data. Then, the proposed model, BIN-PU, generates pseudo labels from the truly positive samples and unlabeled data. Finally, existing backbone CPI prediction models are trained with the generated pseudo positive/negative labels and truly positive samples, along with a new proposed, weighted positive loss function for the PU learning. The details are as follows.

**Fig. 5.**
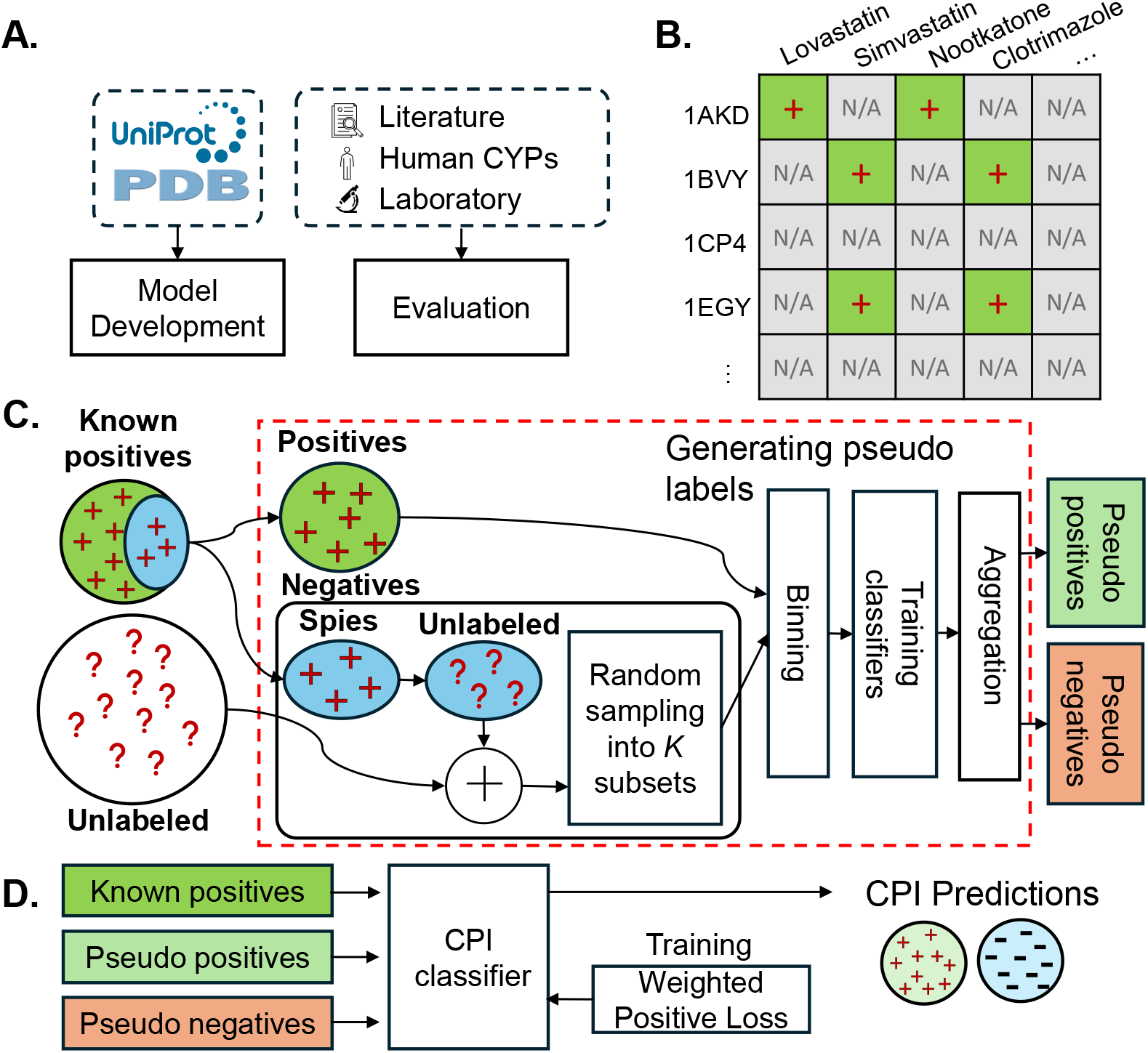
Overview of the proposed framework. (A) We collected datasets from Swiss-Prot and Protein Data Bank (PDB) for model development and assessed the model using literature, human data, and laboratory results for model evaluation. (B) Unlabeled data is generated from known positive samples. (C) A binning-based BIN-PU strategy generates pseudo labels from multiple classifiers trained on truly positive and unlabeled data. (D) Finally, the CPI prediction models are trained with an adaptive loss function using these pseudo labels along with known positive samples.

#### 4.2.1 Generating pseudo labels from positive only data

Pseudo positive and negative labels are computationally annotated by BIN-PU. First, pairwise combinations between unique proteins and compounds in truly positive interactions generate unlabeled samples, where truly positive interactions are excluded. The large volume of the unlabeled samples include both likely positive and negative interactions. Multiple bins are created by sampling from the positive and unlabeled interaction data. Due to the scarcity of positive samples and the abundance of unlabeled data, each bin includes a fixed small number of the truly positive samples and large unlabeled data. Each bin trains a binary CPI classifier, where unlabeled data is considered as negative for the model training. Then, unlabeled samples are categorized into pseudo positives and negatives by computing the average prediction scores of the multiple CPI classifiers.

Specifically, given *N* numbers of known positive training samples, 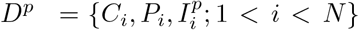, where *C*_*i*_ is a SMILE representation of a compound, *P*_*i*_ is a protein sequence, and 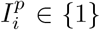 indicates the interaction between the proteins and compounds from all positives in the given dataset. Unlabeled samples, denoted as 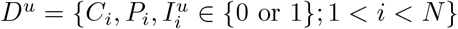, are generated by the pairwise combinations of *C*_*i*_ and *P*_*i*_, excluding known interactions, *I*_*i*_. The positive dataset *D*^*p*^ is split into two: positive samples for training 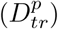 and spy samples for validation 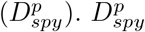 are added to an unlabeled set, i.e., 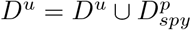, which is used for the validation set to assess the BIN-PU’s performance. For training CPI prediction model, 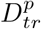 and *D*^*u*^ are utilized as positive and negative samples respectively for training, and 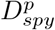 for validating the model.

Let *K* be the number of bins and *f* (·) be an existing CPI prediction model that requires positive and negative interaction samples to train. The negative set *D*^*u*^ is randomly sampled into *K* subsets, 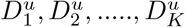, where each 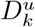 represents a subset of *D*^*u*^ for the *k*-th bin. The training set, *D*_*k*_, for the *k*-th bin is then created by combining 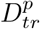 with the *k*-th subset of *D*^*u*^, as:

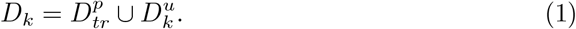

The average posterior probability across all bins are computed as follows:

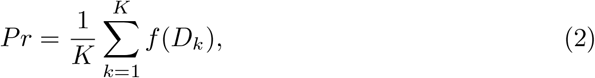

where *f* (*D*_*k*_) is a CPI model with the training set (*D*_*k*_) obtained from bin *k*, and 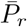 is the averaged probability score across all *K* bins. Pseudo scores 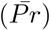 are generated by the averaged probability score. The top-ranked interactions (e.g., 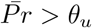) are labeled as pseudo-positives (Ψ_*pos*_), whereas the bottom-ranked samples (e.g., 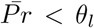) are labeled as pseudo-negatives (Ψ_*neg*_). *θ* is a hyper-parameter that determines reliable pseudo-labels. We set *θ*_*u*_ = 0.8 and *θ*_*l*_ = 0.2 in this study. The remains are unlabeled, which are not used for further training.

#### 4.2.2 Spies capture rate

During the generation of pseudo labels, a spies capture rate assesses the model’s performance of each bin in identifying the known positives of ‘spies’ from unlabeled dataset. These spies are initially parts of the known positive samples but are set aside for the dual purpose: to contribute to training the model to distinguish known positives from unlabeled sets and to evaluate the model’s capability to correctly identify them as positives. The spies capture rate can be computed by:

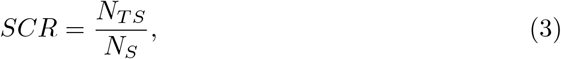

where *N*_*TS*_ represents the number of spies predicted as positives, and *N*_*S*_ represents the total number of spies. A higher SCR (e.g., close to one) indicates that the model is effective at identifying positive pseudo labels from negatives. However, a limitation of this metric arises when the model incorrectly predicts a large number of unlabeled samples as positive. In such cases, the SCR may appear artificially high, because the model would correctly identify most spy samples as positives. However, labeling the top *θ*_*u*_ and bottom *θ*_*l*_ as pseudo-positives and negatives respectively prevent the limitation in our strategy.

#### 4.2.3 Weighted positive loss function for PU learning

Along with the labeled samples of pseudo-samples and known positive interactions, a CPI model is trained with adaptive weights. Traditional loss functions (e.g., binary cross entropy) typically minimize classification errors, considering the same weights of the misclassifications in the truly positive samples and pseudo positive samples. To tackle this limitation, we propose a weighted positive loss term that weighs more to known positives for PU learning, ensuring that the model reduces the misclassification to truly positive samples. The proposed loss function consists of two terms, Binary Cross Entropy (BCE) and the weighted positive loss. The weighted positive loss term penalizes the errors of the predictions in truly positive instances, while the BCE term trains the model using entire samples as a binary classifier. The incorporation of the weighted positive loss term with BCE enhances the calibration of the predicted probabilities in the PU learning setting. The proposed loss function is formulated as:

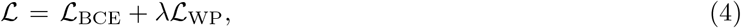

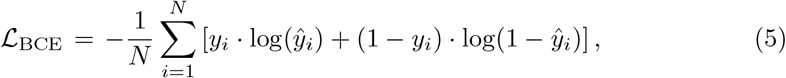

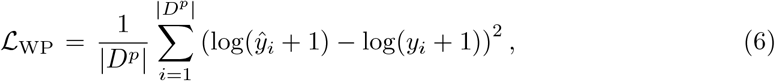

where *N* represents the combination of truly positive, pseudo positive and negative samples (i.e., *N* = |*D*^*p*^| + |Ψ_pos_ |+ |Ψ_neg_ |), *y*_*i*_ is the true label for instance *i, ŷ*_*i*_ is the predicted label of instance *i*, and *λ* represents a weight coefficient assigned to the positive class during the model training.

## Acknowledgements

This research was supported by the National Science Foundation Major Research Instrumentation (NSF MRI) (Grant#:2117941), the Institute of Information & Communications Technology Planning & Evaluation (IITP) grant funded by the Korean government (MSIT) (No. 2021-0-01581). Additional support was provided by the Bio & Medical Technology Development Program (RS-2024-00441423) and the Basic Science Research Program (RS-2023-00276255), both funded by the National Research Foundation of Korea (NRF).

## Availability of data and code

https://github.com/datax-lab/cyp

