## Supplementary for "Prediction of bacterial protein-compound interactions with only positive samples"

### Supplementary Note S1. Dataset gathering criteria for model development

We defined the criteria for data inclusion and exclusion for the data gathering. Keywords for inclusion included “LANOSTEROL 14-ALPHA-DEMETHYLASE”, “P450 MONOOXYGENASE”, and “P450 epoxidase” to capture relevant enzymatic activities and interactions. In contrast, exclusion keywords included “cyclophilin” and “Pyruvate dehydrogenase” to avoid non-relevant proteins, ensuring focus on true CYP interactions. The complete list of keywords is in Table S1. Additionally, we analyzed ligand characteristics, including carbon number and functional type, to filter out inactive compounds or those located outside the active site. We excluded ligands marked as buffers, cofactors, or irrelevant small molecules (e.g., DMSO or buffer ions). This data refinement is shown in Table S2, which categorizes ligands by their inclusion criteria.

**Table S1. Protein names that are included or excluded based on keyword search results in the PDB database.**

| | Keywords |
|---|---|
| Keywords for inclusion | “LANOSTEROL 14-ALPHA-DEMETHYLASE”, “P450 MONOOXYGENASE”, “PROSTACYCLIN SYNTHASE”, ”P450 epoxidase”, “STEROL 14ALPHA-DEMETHYLASE”, ”P450cin”, “ALLENE OXIDE SYNTHASE”, “6-DEOXYERYTHRONOLIDE B HYDROXYLASE”, “PROSTAGLANDIN I SYNTHASE”, “putative cytochrome P450”, “STEROL 14-ALPHA-DEMETHYLASE”, “CAMPHOR 5-MONOOXYGENASE”, “LANOSTEROL 14-ALPHA DEMETHYLASE”, “PIMD PROTEIN”, “STEROL 14 ALPHA-DEMETHYLASE”, ”P-450-LIKE PROTEIN”, “1,5-DIHYDROXYVITAMIN D(3) 4-HYDROXYLASE”, “P450 HEME-THIOLATE PROTEIN”, “STEROL 14-ALPHA DEMETHYLASE”, ”Vitamin D hydroxylase”, “Cholesterol 24-hydroxylase”, “Fatty acid alpha-hydroxylase”, “CHOLESTEROL 4-HYDROXYLASE”, “PUTATIVE MONOOXYGENASE”, “CHOLESTEROL SIDE-CHAIN CLEAVAGE ENZYME”, “TERMINAL OLEFIN-FORMING FATTY ACID DECARBOXYLASE”, “STEROID 1-HYDROXYLASE”, “P450-like protein”, “STEROID 17-ALPHA-HYDROXYLASE/17,0 LYASE”, “PENTALENIC ACID SYNTHASE”, “CHOLESTEROL 7-ALPHA-MONOOXYGENASE”, “Cytochrome P-450”, “CYP17A1”, “MYCOCYCLOSIN SYNTHASE”, “STEROL 14-DEMETHYLASE”, “PENTALENOLACTONE SYNTHASE”, “AROMATASE”, “P450-LIKE ENZYME”, “FERRUGINOL SYNTHASE”, “Fatty-acid peroxygenase”, “PROTEIN LUTEIN DEFICIENT 5”, “METHYL-BRANCHED LIPID OMEGA-HYDROXYLASE”, “HYPOTHETICAL CYTOCHROME P450”, “Aromatic O-demethylase”, “CYTOCHROME P450 BM-3”, “SalCYP” |

| | |
|---|---|
| Keywords for exclusion | “cyclophilin”, “CYTOCHROME P460”, “GREEN FLUORESCENT PROTEIN”, “Pyruvate dehydrogenase”, “NUCLEAR RECEPTOR SUBFAMILY 5 GROUP A MEMBER”, “ferredoxin”, “Histone H3.1”, “AA3-600 quinol oxidase subunit I”, “CYTOCHROME-C55”, “HEMOGLOBIN” |

**Table S2. Ligand characteristics used for dataset refinement**

| Ligand ID | Carbon number | Characteristic |
|---|---|---|
| TRS | C4 | No substrate |
| SRT | C4 | No substrate |
| MLA | C3 | No substrate |
| GOL | C3 | Buffer |
| SPK | C10 | No substrate |
| MES | C6 | Buffer |
| PEG | C4 | Buffer |
| XE | C0 | No substrate |
| BME | C2 | No substrate |
| DTT | C4 | No substrate |
| IPA | C3 | No substrate |
| DMS | C2 | Solvent |
| BEN | C7 | No substrate |
| SIN | C4 | Buffer |
| IMD | C3 | Buffer |
| PG0 | C5 | Buffer |
| MRD | C6 | Buffer |
| EPE | C8 | Buffer |
| MPD | C6 | No substrate |
| TLA | C4 | No substrate |
| FMT | C1 | Small molecule |
| PGE | C6 | No substrate |
| MTN | C10 | Solvent |
| PG4 | C8 | Buffer |
| 1T4 | C18 | No substrate |
| MSE | C5 | Linker |
| CSD | C3 | Linker |
| CME | C5 | Linker |

| | | |
|---|---|---|
| CSO | C3 | Linker |
| 1PE | C10 | Located outside the active site |
| 5KK | C18 | No substrate |
| SPD | C7 | No substrate |
| RU8 | C34 | Photosensitizer |
| TAM | C7 | No substrate |
| DIO | C4 | No substrate |
| BCN | C6 | Located outside the active site |
| MPO | C7 | Buffer |
| SER | C3 | Located outside the active site |
| SPM | C10 | Located outside the active site |
| 17Q | C19 | Located outside the active site |
| OXY | C0 | Small molecule |
| HOA | C0 | Small molecule |
| NO | C0 | Small molecule |
| CMO | C1 | Small molecule |
| FMN | C17 | Cofactor |
| FES | C0 | Cofactor |
| MNR | C34 | Cofactor |
| HTG | C13 | Cofactor |
| 1N0 | C14 | Cofactor |

### Supplementary Note S2. Model implementation

We implemented the three CPI backbone models, including CPIprediction, TransformerCPI, and SSNet. For the preprocessing, we followed their proposed encoding schemes for proteins and compounds. Specifically, for CPIprediction, we encoded protein sequences using overlapping n-grams (3-mers). Each n-gram was converted into a trainable embedding vector of dimension 128, and the embedding was processed through stacked convolutional layers, followed by max-pooling to extract local sequence motifs. For compound encoding, we converted SMILES strings into molecular graphs using RDKit. Each atom was encoded as a feature vector that included atom type, hybridization, aromaticity, formal charge, degree, and number of hydrogens, while bonds were encoded in an adjacency matrix specifying bond types. These molecular graphs were processed using a graph neural network to generate compound-level graph embeddings. For TransformerCPI, we trained a Word2Vec model on the protein sequences from the training dataset to learn task-specific embeddings for amino acids. Each amino acid was mapped to its corresponding dense vector, and the sequence of embeddings was processed through transformer encoder layers. For compound encoding, we computed 34-dimensional molecular descriptors for each compound using RDKit. These descriptors summarize atom types, hybridization, valence, aromaticity, and bond connectivity. We fed the descriptor vectors into a feedforward encoder and processed them together with the protein embeddings using attention layers. For SSNet, we computed structural encodings from protein 3D conformations. We extracted C-α atom coordinates from the PDB files, and curvature ($\kappa$) and torsion ($\tau$) values were calculated for each residue to generate a two-channel (2×sequence-length) representation. We processed these $\kappa$–$\tau$ sequences using multi-branch one-dimensional convolutional neural networks with kernel sizes of 5, 10, 15, and 20, followed by global max-pooling. For compound encoding, we constructed molecular graphs from SMILES strings using RDKit, following the same atom and bond feature extraction procedure as done in CPIprediction. We additionally computed molecular fingerprints, including ECFP-like circular fingerprints. We concatenated molecular fingerprints with the graph-derived features to form hybrid dense vectors, which we passed through fully connected layers to generate compound embeddings compatible with the protein $\kappa$–$\tau$ representations.

For BIN-PU, we followed the procedure illustrated in Fig. 2. We first generated unlabeled data by creating pairwise combinations of proteins and compounds, excluding known positive interactions. We randomly divided the unlabeled data into $K$ subsets (bins) and trained multiple CPI classifiers, each using a combination of truly positive samples and one subset of unlabeled samples for each bin. We aggregated the posterior probabilities from these classifiers to assign pseudo labels, selecting high-confidence samples as pseudo positives and pseudo negatives. We then trained the CPI backbone models (CPIprediction, TransformerCPI, and SSNet) in a fully supervised manner using truly positive, pseudo positive, and pseudo negative samples together. During training, we applied a weighted positive loss function to give higher importance to truly positive samples and to minimize misclassification of biologically relevant interactions.

For PUCPI, we implemented the training as described in the original study. All unlabeled samples were treated as negatives. We utilized the same datasets, data splits, and model configurations as in BIN-PU to ensure a fair comparison. We trained PUCPI coupled with the CPI backbone models directly with truly positive and unlabeled (assumed negative) samples. PUCPI did not include any pseudo-label generation, however, we used the pseudo-labels generated by BIN-PU for evaluating the performance of PUCPI. We also considered the original PUCPI using a biased-SVM without experimentally validated negative samples. PUCPI was applied to human enzyme data by utilizing substructures and the PFAM database for data preprocessing. However, due to the unavailability of the database for bacterial CYP data, we slightly modified the training setting of PUCPI. We performed the same data preprocessing of CPIPrediction for PUCPI. Specifically, we first extracted feature vectors for protein and compound vectors. The compound vector was multiplied with an adjacency vector to include bond information. We fed the multiplied vector to a 1D CNN to match the number of output channels as a protein vector. We computed the tensor product between the protein vector and the converted compound vector. Following that, a biased-SVM classifier was trained on the tensor product by providing more weight to known positives. The hyperparameters, including *c* (tradeoff between training error and margin) and *j* (cost factor) of biased-SVM loss function, were tuned using the validation data. We used the same training, validation, and test data for a fair comparison.

### Supplementary Note S3. Hyperparameter tuning for BIN-PU

We obtained the optimal hyperparameters, including the bin size and the lambda value for the weighted positive loss function, that optimized the performance on the validation data for each experiment. Specifically, the optimal number of bins (bin size) was obtained to maximize the Spies Capture Rate (SCR). We evaluated bin sizes of 10, 20, and 30 across the three CPI backbone models and selected 20 as the optimal bin size, as it consistently achieved the highest SCR (Table S3, Fig. S1A).

We also tuned the lambda parameter in the weighted positive loss function to control the weight assigned to truly positive samples. We tested lambda values of 1, 10, 100, and 1000 across the three CPI backbone models and selected lambda = 100 as the optimal value, as it consistently resulted in the highest F1-scores across experiments (Table S4, Fig. S1B).

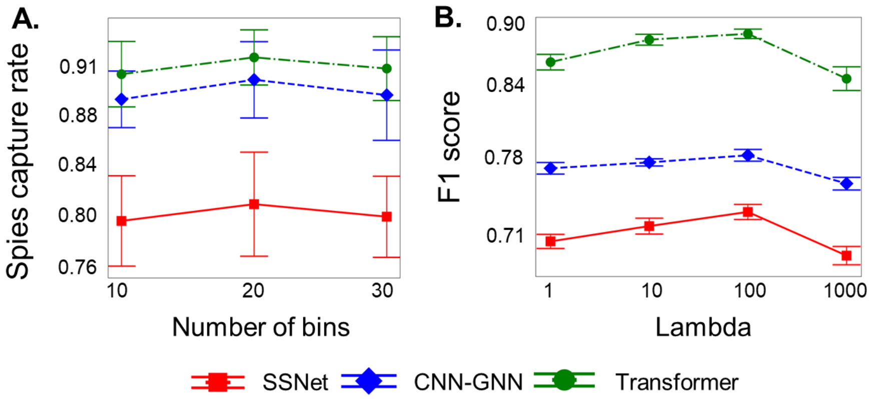


**Figure S1. Comparison of BIN-PU performance across varying parameters using error bar plots**. (A) Spies capture rate across multiple bins, (B) Lambda values vs F1 score during training of CPI model with weighted positive loss function, and the error bars represent the standard deviation across 10 iterations for each model.

**Table S3. Performance comparison varying bin sizes**

| Model | 10 bins | 20 bins | 30 bins |
|---|---|---|---|
| CPIprediction | 0.891 ± 0.00031 | **0.904 ± 0.001664** | 0.886 ± 0.00142 |
| TransformerCPI | 0.908 ± 0.00224 | **0.920 ± 0.00144** | 0.914 ± 0.00096 |
| SSNet | 0.81 ± 0.0052 | **0.846 ± 0.0064** | 0.832 ± 0.00664 |

**Table S4. Performance comparison of CPI backbone model with varying lambdas**

| Model | lambda=1 | lambda =10 | lambda =100 | lambda =1000 |
|---|---|---|---|---|
| CPIprediction | 0.766 ± 0.00333 | 0.77 ± 0.00563 | **0.774 ± 0.00166** | 0.752 ± 0.00113 |
| TransformerCPI | 0.854 ± 0.00782 | 0.872 ± 0.00056 | **0.876 ± 0.00176** | 0.838 ± 0.00068 |
| SSNet | 0.704 ± 0.00146 | 0.718 ± 0.00335 | **0.728 ± 0.00332** | 0.686 ± 0.00889 |

**Table S5. List of CYPs included in literature papers**

| CYP name | PDB ID | Reference |
|---|---|---|
| *Ba*CYP106A2 | 5XNT | Kim, K. H., Lee, C. W., Dangi, B., Park, S. H., Park, H., Oh, T. J., & Lee, J. H. (2017). Crystal structure and functional characterization of a cytochrome P450 (BaCYP106A2) from Bacillus sp. PAMC 23377. Journal of Microbiology and Biotechnology, 27(8), 1472-1482. |
| CYP106A2 | 4YT3 | Schmitz, D., Janocha, S., Kiss, F. M., & Bernhardt, R. (2018). CYP106A2—a versatile biocatalyst with high potential for biotechnological production of selectively hydroxylated steroid and terpenoid compounds. Biochimica et Biophysica Acta (BBA)-Proteins and Proteomics, 1866(1), 11-22. |
| CYP109E1 | 5L91 | 1. Jóźwik, I. K., Kiss, F. M., Gricman, Ł., Abdulmughni, A., Brill, E., Zapp, J., ... & Thunnissen, A. M. W. (2016). Structural basis of steroid binding and oxidation by the cytochrome P450 CYP 109E1 from Bacillus megaterium. The FEBS Journal, 283(22), 4128-4148.<br>2. Putkaradze, N., Litzenburger, M., Hutter, M. C., & Bernhardt, R. (2019). CYP109E1 from Bacillus megaterium Acts as a 24-and 25-Hydroxylase for Cholesterol. ChemBioChem, 20(5), 655-658.<br>3. Abdulmughni, A., Jóźwik, I. K., Putkaradze, N., Brill, E., Zapp, J., Thunnissen, A. M. W., ... & Bernhardt, R. (2017). Characterization of cytochrome P450 CYP109E1 from Bacillus megaterium as a novel vitamin D3 hydroxylase. Journal of Biotechnology, 243, 38-47. |

| | | |
|---|---|---|
| | | 4. Putkaradze, N., König, L., Kattner, L., Hutter, M. C., & Bernhardt, R. (2020). Highly regio-and stereoselective hydroxylation of vitamin D2 by CYP109E1. Biochemical and biophysical research communications, 524(2), 295-300.<br>5. Putkaradze, N., Litzenburger, M., Abdulmughni, A., Milhim, M., Brill, E., Hannemann, F., & Bernhardt, R. (2017). CYP109E1 is a novel versatile statin and terpene oxidase from Bacillus megaterium. Applied Microbiology and Biotechnology, 101, 8379-8393. |
| CYP109B1 | 4RM4 | Girhard, M., Klaus, T., Khatri, Y., Bernhardt, R., & Urlacher, V. B. (2010). Characterization of the versatile monooxygenase CYP109B1 from Bacillus subtilis. Applied microbiology and biotechnology, 87, 595-607. |
| *Ba*CYP106A6 | 8HG9 | Kim, K. H., Do, H., Lee, C. W., Subedi, P., Choi, M., Nam, Y., ... & Oh, T. J. (2022). Crystal structure and biochemical analysis of a cytochrome P450 steroid hydroxylase (BaCYP106A6) from Bacillus species. Journal of Microbiology and Biotechnology, 33(3), 387. |

### Supplementary Note S4. Biological experiment with uncurated bacterial enzymes

Dr. Nelson named CYP154C9, CYP107G9, CYP1047A9, CYP106A4, and CYP106A5 through sequence blast (see Table S6). Each CYP was inserted in the pET28a vector and introduced into Escherichia coli C41(DE3). CYP was incubated in Luria–Bertani (LB) broth supplemented with antibiotics (100 µg/ml of ampicillin) at 37°C. A seed culture of CYP was grown in LB and maintained until an optical density at 600nm (OD600) of 0.6-0.8 was reached. 0.5 mM of FeCl3·6H2O and 1 mM of 5-aminolevulinic acid hydrochloride (5-ALA) were added to assist in heme formation. After cooling to 20°C, CYP was overexpressed by injecting 0.5 mM IPTG and incubated for 3 days. The harvested cell was suspended in a 50 mM potassium buffer (pH 7.4). The cell extract was sonicated to obtain the soluble protein solution. The protein solution was mixed with His-tag resin solution and eluted with 20-, 100-, and 250-mM imidazole potassium phosphate buffer. The eluted fraction containing the protein was concentrated using an Ultra centrifugal filter (Millipore, Ireland).

For the *in vitro* assay, steroid substrates (4-androstenedione, progesterone, and nandrolone) were used. A stock solution (100 mM) of the substrate was dissolved in DMSO and stored until use. The *in vitro* reaction was carried out in a total volume of 250 µl in a 50-mM potassium buffer (pH 7.4) consisting of 3 µM CYP, 2.5 mM iodo-benzene, and 200 µM steroid substrates. The CYP154C9, CYP107G9, and CYP1047A9 used iodo-benzene for the electron transfer system. The 10- µM CYP106A4 and A5 reacted with 100- µM FdX, 0.1U FdR, 200- µM steroid, 5- mM $MgCl_2$, 100- µg/ml catalase, and NADH generation system (1- mM NADH, 1- U glucose-6-phosphate dehydrogenase (G6P-DH), and 10- mM glucose-6-phosphate) in a 50- mM potassium buffer (pH 7.4). The reaction was initiated by adding 1 mM of NADH. Reaction mixtures were incubated at 30°C for 2 h, extracted twice with an equal volume of ethyl acetate, and dried completely. The dried samples were dissolved in methanol for HPLC analysis. HPLC analysis conditions were described previously [1].

**Table S6. Strain of microorganisms and the CYP family for each strain of CYP used in this study**

| CYP Name | Species | Strain or isolated area |
|---|---|---|
| CYP154C9 | Streptomyces alboniger | KCCM[a] 12598 |
| CYP1047A9 | Streptomyces sp. | KCCM 40463 |
| CYP107G9 | Actinomycete sp. | ATCC[b] 53650 |
| CYP106A4 | Paenibacillus sp. | Antarctica |
| CYP106A5 | Paenibacillus sp. | Antarctica |

[a]KCCM: Korean Culture Center of Microorganisms

[b]ATCC: American Type Microbiology

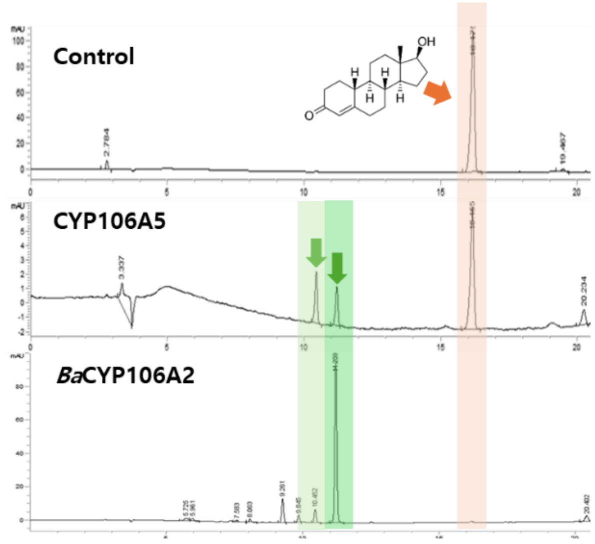


**Figure S2**. **HPLC chromatogram for comparison of *in vitro* assay results.** It used CYP106A5 and *Ba*CYP106A2 as target proteins and nandrolone as a ligand compound [1]. The orange arrows and boxes represent the nandrolone peaks, and the two green arrows and boxes represent the products, respectively. The main product of *Ba*CYP106A2 is 15β hydroxy compound.

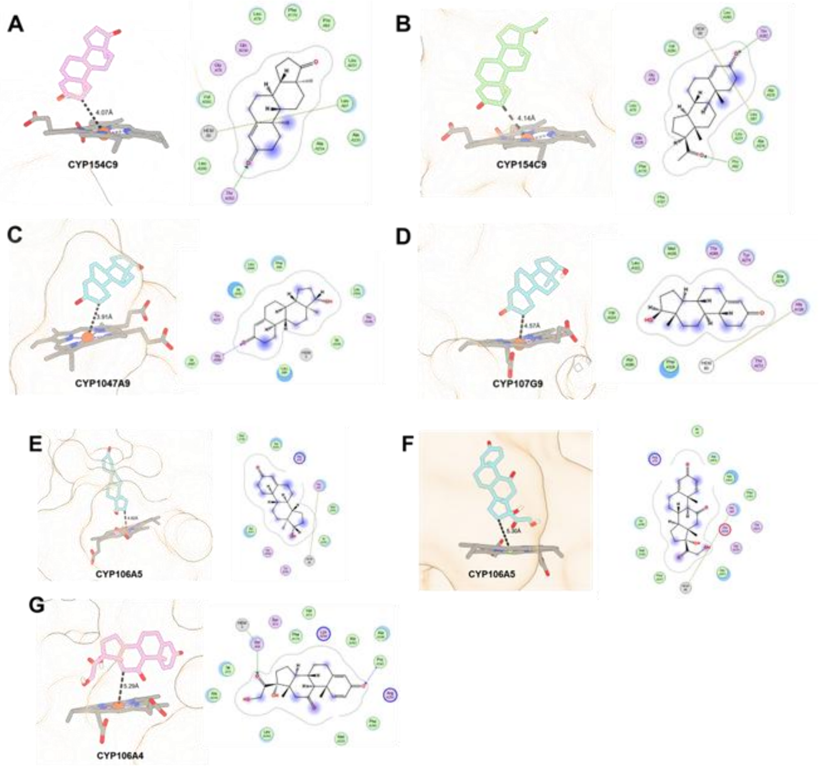


**Figure. S3**. Binding poses in target protein and 2D molecular docking diagrams showing the binding interactions of the best docked ligand compounds 4-androstenedione (A), progesterone (B), nandrolone (C, D and E), and prednisone (F and G).

### Supplementary Note S5. Biophysical experiment with uncurated bacterial enzymes

We also performed molecular docking to further assess the potential protein-ligand interactions. Since the three-dimensional structures of cytochrome P450 (CYP) protein strains are not available in the Protein Data Bank, the Chai Discovery web server [2] was used to generate homology models for all CYP isoforms. The pTM scores of all the homology models, along with the corresponding amino acid sequence lengths, are provided in Table S7. All ligand structures were downloaded from the PubChem database in SDF format and converted to PDBQT format by using Meeko v0.6.1 Python tool [https://github.com/forlilab/Meeko]. The grid box was centered on the heme iron of the CYP proteins, with a dimension of 18 × 18 × 18 Å. Molecular docking was performed using AutoDock Vina version 1.2.3 [3,4]. The docking scores and the distance between the heme iron and the nearest carbon atom are reported in Table 1. Snapshots of the protein-ligand docking were captured using ChimeraX [5], and ligand-protein interactions were analyzed by using Molecular Operating Environment (MOE) 2024.06 [6].

**Table S7. Homology model (*pTM*) scores and amino acid lengths of the CYP Isoforms**

| CYP Isoforms | pTM score | Amino acid length |
|---|---|---|
| CYP154C9 | 0.9616 | 404 |
| CYP1047A9 | 0.9417 | 455 |
| CYP107G9 | 0.9163 | 439 |
| CYP106A4 | 0.9592 | 410 |
| CYP106A5 | 0.9333 | 418 |
